# The Effect of Productive HPV16 Infection on Global Gene Expression of Cervical Epithelium

**DOI:** 10.1101/295402

**Authors:** Sa Do Kang, Sreejata Chatterjee, Samina Alam, Anna C. Salzberg, Janice Milici, Sjoerd H. van der Burg, Craig Meyers

## Abstract

Human papillomavirus (HPV) infection is the world’s most common sexually transmitted infection, and is responsible for most cases of cervical cancer. Previous studies of global gene expression changes induced by HPV infection have focused on the cancerous stages of infection, and therefore, not much is known about global gene expression changes at early pre-neoplastic stages of infection. We show for the first time, global gene expression changes of early stage HPV16 infection in cervical tissue using 3-dimensional organotypic raft cultures that produce high levels of progeny virions.

cDNA microarray analysis showed that a total of 594 genes were upregulated and 651 genes were downregulated at least 1.5-fold with HPV16 infection. Gene ontology analysis showed that biological processes including cell cycle progression and DNA metabolism were upregulated, while skin development, immune response, and cell death were downregulated with HPV16 infection in cervical keratinocytes. Individual genes were selected for validation at the transcriptional and translational levels including UBC, which was central to the protein association network of immune response genes, and top downregulated genes RPTN, SERPINB4, KRT23, and KLK8. In particular, KLK8 and SERPINB4 have shown to be upregulated in cancer, which contrasts our results.

Organotypic raft cultures that allow full progression of the HPV life-cycle have allowed us to identify novel gene modulations and potential therapeutic targets of early stage HPV infection in cervical tissue. Additionally, our results suggest that early stage productive infection and cancerous stages of infection are distinct disease states expressing different transcriptomes.

**Importance:** Persistent HPV infection is responsible for most cases of cervical cancer. Transition from precancerous to cancerous stages of HPV infection is marked by a significant reduction in virus production. Most global gene expression studies of HPV infection have focused on the cancerous stages. Therefore, little is known about global gene expression changes at precancerous stages. For the first time, we measured global gene expression changes at precancerous stages of HPV16 infection in human cervical tissue producing high levels of virus. We identified a group of genes that are typically overexpressed in cancerous stages to be significantly downregulated at the precancerous stage. Moreover, we identified significantly modulated genes that have not yet been studied in the context of HPV infection. Studying the role of these genes in HPV infection will help us understand what drives the transition from precancerous to cancerous stages, and may lead to development of new therapeutic targets.

## Introduction

Human papilloma virus (HPV) infection is the world’s most common sexually transmitted infection with approximately 291 million women worldwide infected with the virus at any given point in time (1). While low-risk HPV types cause benign warts in the anogenital area, persistent infection with high-risk HPV types can give rise to various cancers of epithelial origin. In particular, HPV is responsible for most cases of cervical cancer, which is the third most common cancer in women worldwide and the most common cancer in women in developing countries (2). For this reason, the mechanism of HPV infection and HPV-mediated oncogenesis has been extensively studied. Early viral proteins E6 and E7 have been identified as oncoproteins that play a critical role in tumorigenesis and tumor maintenance by inhibiting tumor suppressor genes *TP53* and *RB1*, respectively (3–10).

HPV initially infects dividing cells of the basal layer of the epidermis via microabrasions. Most infections are naturally resolved, but in some cases, the virus establishes a persistent infection, which may subsequently progress to precancerous lesions that are histologically graded as cervical intraepithelial neoplasia (CIN) I to III. When left untreated, these neoplastic lesions may ultimately develop into carcinoma in situ and invasive cancer. In persistently infected cervical tissue with normal or low-grade dysplasia the HPV genome is maintained episomally and infectious viral particles are produced. In contrast, progression to severe dysplasia and invasive cancer is marked by integration of viral genome into the host genome that typically results in disruption of the E2 gene, subsequent upregulation of oncogenes E6 and E7, and abrogation of virion production (11, 12).

In the past, many studies have looked into changes in whole genome expression profiles of precancerous and cancerous lesions in order to better understand the progression of persistent HPV infection to cervical cancer. However, most of these studies focus on neoplastic lesions and cancerous lesions (13–21) and, therefore, there is a gap in knowledge of global gene expression in earlier pre-neoplastic stages of the disease when HPV establishes productive infection in the host. In two recent studies, human keratinocytes persistently infected with HPV16 were used to measure global gene expression changes (22, 23), but they used keratinocytes derived from foreskin, which may not be appropriate for modeling infection in cervical tissue since HPV may have tissue-specific effects (24). Furthermore, these studies used monolayer cell cultures that do not produce mature virions by disallowing the virus to progress through its differentiation-dependent replication life-cycle (25, 26). In other studies, overexpression tools were used to examine the effect of specific HPV oncoproteins on global gene expression (27, 28). Since these studies only examine the effect of individual viral proteins, they do not account for the full picture of HPV infection in the natural environment.

In this study, we used oligonucleotide microarrays to measure global gene expression changes in early passage HPV16-infected human cervical keratinocytes (16HCK). Organotypic raft cultures were used in order to allow the virus to go through its full life-cycle. A total of 594 and 651 genes were at least 1.5-fold upregulated and downregulated, respectively. Gene ontology analysis of upregulated genes identified biological processes that were significantly represented including the cell cycle process and DNA metabolism. In contrast, biological processes that were significantly represented with downregulated genes included epidermis development, extracellular matrix disassembly, and regulation of NF-κB signaling.

## Results

### Microarray analysis of HPV16 infection in cervical tissue

Global gene expression changes in cervical tissue infected with HPV16 at a pre-neoplastic state were measured by conducting cDNA microarray analysis (GEO accession no. GSE109039) on 10-day organotypic raft tissue from early passage human cervical keratinocytes (HCK) persistently infected with HPV16 (16HCK). In order to create 16HCK cell lines, human cervical tissue from biopsies were acquired, processed, and cultured with keratinocyte-selective medium to grow primary HCKs. The HCKs were then electroporated with the HPV16 genome to establish persistent infection and immortalization. Since HPV replication is differentiation-dependent, monolayer cell culture systems cannot produce HPV virions and provide limited insight into the effect of productive infection that happens *in vivo*. Also, it is difficult to study early stages of HPV infection in clinical samples because early stages of infection are largely asymptomatic and patients typically present at a neoplastic or cancerous stage. Moreover, clinical samples may be infected with any of the numerous types and variants of HPV making it difficult to study the effect of a specific virus in a controlled environment. Therefore, we used organotypic raft cultures as previously described, which allows us to observe the full HPV life-cycle in 3-dimensional (3D) tissue at a precancerous stage of infection when HPV particles are maximally produced. The organotypic raft tissue was harvested at 10 days of culture for microarray analysis when particle production is most active, and 20 days of culture when particle accumulation is at its maximum to measure the viral titer showing that virus particles are produced at high levels (Table 1). By infecting each cell line with the same virus and subjecting the raft cultures to the same condition, we are able to minimize variables and observe the specific effect of HPV infection on cervical tissue.

**Table 1.** Viral titers of organotypic rafts

| Raft Identification | Viral particles per raft |
| --- | --- |
| 16HCK-1, replicate 1 | $9.38 \times 10^7$ |
| 16HCK-1, replicate 2 | $1.18 \times 10^7$ |
| 16HCK-2, replicate 1 | $5.25 \times 10^8$ |
| 16HCK-2, replicate 2 | $6.5 \times 10^9$ |
| 16HCK-3, replicate 1 | $5.81 \times 10^7$ |
| 16HCK-3, replicate 2 | $1.6 \times 10^9$ |

The experiment was conducted in three individually derived 16HCK cell lines in duplicates and raft tissue from uninfected HCKs served as control. Out of the 34,575 genes that were analyzed with cDNA microarray, a total of 1,245 genes were modified at least 1.5-fold (p<0.05) and 533 genes were modified at least 2-fold (p<0.05) with HPV16 infection as compared to uninfected control (Table S1). Of those genes that were significantly modulated at least 1.5-fold, 594 genes were upregulated and 651 genes were downregulated. Table 2 and 3 show the 50 most upregulated and downregulated genes in the microarray analysis. Amongst the top 50 upregulated genes are those associated with cell cycle (*CDKN2A*, *CDC7*, *NASP*, *MDC1*, *NFIX*, *FOXQ1*). In contrast, the 50 most downregulated genes span from those involved in differentiation (*RPTN*, *LCE1D*, *LCE3C*, *LCE1E*, *S100A7*), ECM-modulation (*KLK6, 8, 10, 13* and *MMP 9, 10*), immune regulation (*LCN2*, *SPNS2*, *FAM3D*, *IL1RN*, *PSG4*, *IL1F7*) to antimicrobial response (*RNASE7*, *PRSS3*, *PRSS2*). This suggests that HPV infection is driving the cell cycle, and disrupting epidermal differentiation and ECM homeostasis while evading the immune and antimicrobial responses by downregulating them.

**Table 2.** Top 50 upregulated genes with productive HPV16 infection

| GENE | FUNCTION | FOLD<br>CHANGE |
| --- | --- | --- |
| *MT1G* | Metallothioneins have a high content of cysteine residues that bind various heavy metals; these proteins are transcriptionally regulated by both heavy metals and glucocorticoids. | 12.032 |
| *MT1H* | Mineral absorption/ metabolism | 7.679 |
| *TGFBR3* | Binds to TGF-beta. Could be involved in capturing and retaining TGF-beta for presentation to the signaling receptors | 7.295 |
| *KLHL35* | Kelch-like family member. Kelch-repeat β-propellers are generally involved in protein-protein interactions | 5.814 |
| *DLK2* | Calcium ion binding and protein dimerization activity | 5.057 |
| *FBLN1* | Cell adhesion/ migration/ ECM architecture organization | 5.056 |
| *ASS1* | Arginine biosynthetic pathway | 4.446 |
| *GPER* | G Protein coupled estrogen receptor; cAMP signaling, calcium mobilization | 4.446 |
| *TDRD9* | Probable ATP binding RNA helicase | 4.393 |
| *TWIST1* | Transcription factor, role in cell lineage determination and differentiation | 4.208 |
| *FANCE* | Member of the Fanconi anemia complementation group E | 4.183 |
| *JAM3* | Cell to cell adhesion in epithelial and endothelial cells | 4.169 |
| *CDKN2A* | Cell cycle regulation | 4.122 |
| *TSPAN4* | Cell surface protein that mediate cell development, | 3.733 |
|  | activation, growth and motility |  |
| *LTBP4* | Involved in assembly, targeting and activation of TGFB1 | 3.718 |
| *ERAP2* | Aminopeptidase involved in peptide trimming, MHC class I presentation | 3.551 |
| *LOC341230* | Similar to argininosuccinate synthetase | 3.454 |
| *MXRA5* | Matrix remodeling associated proteins | 3.281 |
| *RPS23* | Component of the 40S ribosomal subunit | 3.23 |
| *C14orf132* | Putative uncharacterized protein | 3.165 |
| *CRIP2* | Putative transcription factor with two LIM zinc binding domains | 3.148 |
| *ANGPTL2* | Member of the vascular endothelial growth factor family | 3.061 |
| *GOLPH4* | Involved in endosome to Golgi protein trafficking | 3.058 |
| *DPYSL2* | Cytoskeletal remodeling, endocytosis | 3.013 |
| *CDC7* | Checkpoint control kinase critical for G1/S transition | 2.981 |
| *FAM134B* | ER anchored autophagy receptor | 2.947 |
| *NASP* | Histone binding protein with a role in cell division, cell cycle progression and proliferation | 2.885 |
| *HEG1* | Calcium ion binding receptor component, may act | 2.866 |
|  | through stabilization of endothelial cell junctions |  |
| *ZCWPW1* | Zinc finger domain containing protein | 2.8 |
| *MTE* | Metallothioneins have a high content of cysteine residues that bind various heavy metals. | 2.786 |
| *MDC1* | Required for checkpoint mediated cell cycle arrest in response to DNA damage | 2.783 |
| *CDH13* | Calcium dependent cell adhesion proteins | 2.775 |
| *OLFML2A* | Extracellular matrix binding protein | 2.762 |
| *LOC392871* | Undetermined gene ortholog | 2.728 |
| *CYBRD1* | Ferric chelate reductase | 2.726 |
| *RASIP1* | Ras interacting protein required for the formation of vascular structures | 2.721 |
| *LOC729137* | Undetermined gene ortholog | 2.7 |
| *GPNMB* | Type 1 transmembrane glycoprotein | 2.693 |
| *MOBKL2B* | May regulate the activity of kinases | 2.672 |
| *LOC388494* | Undetermined gene ortholog | 2.645 |
| *C10orf54* | Putative uncharacterized protein | 2.644 |
| *E2F2* | E2F family of transcription factors | 2.572 |
| *LFNG* | Transferase enzyme | 2.571 |
| *LOC100133866* | Undetermined gene ortholog | 2.568 |
| *NFIX* | DNA binding protein, capable of activating transcription and replication | 2.56 |
| *P4HTM* | Prolyl hydroxylases | 2.553 |
| *RELL1* | Receptor expressed in lymphoid tissue like 1 | 2.551 |
| *FOXQ1* | FOX genes involved in gene development, cell proliferation and tissue specific gene expression | 2.53 |
| *TJAP1* | Tight junction associated protein | 2.527 |
| *HOXA11AS* | Non-coding RNA gene | 2.525 |
| *FN3KRP* | Phosphorylates psicosamines and ribulosamines | 2.514 |

**Table 3.** Top 50 downregulated genes with productive HPV16 infection

| **GENE** | **FUNCTION** | **FOLD**<br>**CHANGE** |
| --- | --- | --- |
| *RPTN* | Involved in the cornified cell envelope formation.<br>Multifunctional epidermal matrix protein. Reversibly binds calcium. | -57.487 |
| *SERPINB4* | May act as a protease inhibitor to modulate the host immune response against tumor cells | -48.016 |
| *TCN1* | Binds Vit B12 and protects it from the acidic environment of the stomach | -34.735 |
| *CST6* | Active cysteine protease inhibitors. Loss of function is associated with progression of primary tumor to a metastatic phenotype | -27.758 |
| *KLK8* | Serine proteases with diverse physiological function | -25.804 |
| *CEACAM6* | GPI anchored glycoprotein and plays a role in cell adhesion | -24.469 |
| *KLK6* | Serine proteases regulated by steroids | -22.088 |
| *KLK13* | Expression regulated by steroid hormones and important marker for breast cancer | -21.464 |
| *RNASE7* | Pancreatic ribosomal protein; broad spectrum antimicrobial activity against bacteria and fungi | -20.029 |
| *PRSS3* | Digestive protease specialized for the degradation of trypsin inhibitors. In the ileum, may be involved in defensin processing, including DEFA5 | -19.968 |
| *LCE1D* | Precursors of the cornified layers of the stratum corneum | -17.868 |
| *MMP9* | Zinc dependent endopeptidases and major proteases | -17.142 |
|  | involved in the degradation of the ECM |  |
| *FLJ22662* | Weak phospholipase activity. May act as an amidase or peptidase | -14.394 |
| *TMEM45B* | Transmembrane protein 45B | -13.469 |
| *DMKN* | Expressed in differentiated layers of the skin. Upregulated in inflammatory disease | -13.186 |
| *C6orf15* | Uncharacterized protein coding gene | -12.641 |
| *TMEM45A* | Paralog of TMEM45B | -12.629 |
| *PNLIPRP3* | Triglyceride lipase activity | -11.232 |
| *KRT23* | intermediate filament protein | -11.074 |
| *ANXA9* | Calcium dependent phospholipid binding protein; binds ECM proteins | -10.728 |
| *LCE3C* | Precursors of the cornified layers of the stratum corneum | -10.044 |
| *S100A7* | Calcium binding proteins involved in cell cycle progression and cellular differentiation | -9.88 |
| *EPS8L1* | Substrate for the Epidermal growth factor receptor | -9.659 |
| *RORA* | Member of NR1 family of nuclear hormone receptors | -9.136 |
| *LCN2* | Iron trafficking protein involved in apoptosis, innate immunity and renal development | -9.134 |
| *LCE1E* | Precursors of the cornified layers of the stratum corneum | -9.067 |
| *SPNS2* | Sphingolipid transporter with critical function in cardiovascular, immunological and neuronal development | -8.787 |
| *FAM3D* | Related to cytokine activity | -8.734 |
| *KLK7* | Member of the kallikrein family of serine proteases | -8.61 |
| *HMOX1* | Heme oxygenase, an essential enzyme in heme catabolism | -8.548 |
| *RNF39* | Role in early phase of synaptic plasticity | -8.5 |
| *SH3BGRL2* | Protein disulphide oxidoreductase activity | -8.478 |
| *C1orf68* | Uncharacterized protein coding gene | -8.417 |
| *POF1B* | Key role in organization of the epithelial monolayers by regulating the actin cytoskeleton | -8.233 |
| *ATP6V1C2* | Subunit of the peripheral V1 complex of the vacuolar ATPase | -8.232 |
| *MMP10* | Zinc dependent endopeptidases and major proteases involved in the degradation of the ECM | -7.953 |
| *LOC730833* | Undetermined gene ortholog | -7.935 |
| *CAPNS2* | Calcium regulated thiol protease, role in tissue remodeling and signal transduction | -7.64 |
| *KLK10* | Member of the kallikrein family of serine proteases | -7.553 |
| *IL1RN* | Inhibits the activity of Interleukin-1 | -7.499 |
| *ABCG1* | Intracellular lipid transport | -7.408 |
| *THEM5* | Role in mitochondrial fatty acid metabolism | -7.287 |
| *CYP2J2* | Role in arachidonic acid metabolism | -7.259 |
| *PRSS2* | Serine protease involved in defensin processing | -7.239 |
| *LOC645869* | Undetermined gene ortholog | -6.803 |
| *PSG4* | May play a role in regulation of innate immune system | -6.559 |
| *FAM63A* | Role in protein turnover, deubiquitinase activity | -6.52 |
| *MAP2* | Stabilizing/stiffening of microtubules | -6.474 |
| *SLC5A1* | Na<sup>+</sup> /glucose cotransporter | -6.468 |
| *IL1F7* | Interleukin 1 related gene | -6.392 |
| *EPB41L3* | Tumor suppressor that inhibits cell proliferation and promotes apoptosis | -6.356 |

While most previous global gene expression studies have focused on neoplastic and cancerous stages of HPV infection (13–21), two previous studies have modeled pre-neoplastic, early stage HPV infection similar to our study (22, 23). However, most of the modulated genes in the two studies did not overlap with our results. One of these studies reported that 135 genes were modulated with HPV infection, but only 38 (28.1%) of these were modulated in the same direction and 4 of them were modulated in the opposite direction in our microarray analysis (22). In the other study, a total of 966 genes were reported to be modulated with HPV infection, but only 15 (1.6%) of these were modulated in the same direction in our microarray analysis (23). The two major differences between our study and the two previous studies is that we use cervical keratinocytes instead of foreskin keratinocytes, and that we use organotypic raft cultures instead of monolayer cultures that do not allow HPV to complete its life cycle. Using keratinocytes derived from cervical tissue, and organotypic raft cultures that allow the virus to complete its life cycle in 3D tissue enables us to capture the whole picture of an early stage HPV infection in the cervix.

While fold-change of the 50 most upregulated genes ranged between 12.1 to 2.5, that of the 50 most downregulated genes ranged between 57.5 and 6 indicating that a greater degree of gene modulation occurred in the downregulated genes than the upregulated genes. Similarly, when the cutoff for inclusion was increased to at least 5-fold modulation, only 6 genes were upregulated in contrast to 72 genes that were downregulated as shown in Fig. 1. This suggests that HPV16 is disrupting the host’s physiology mostly by dampening many of the normal processes that may interfere with the virus’s survival and replication.

**FIG 1.**
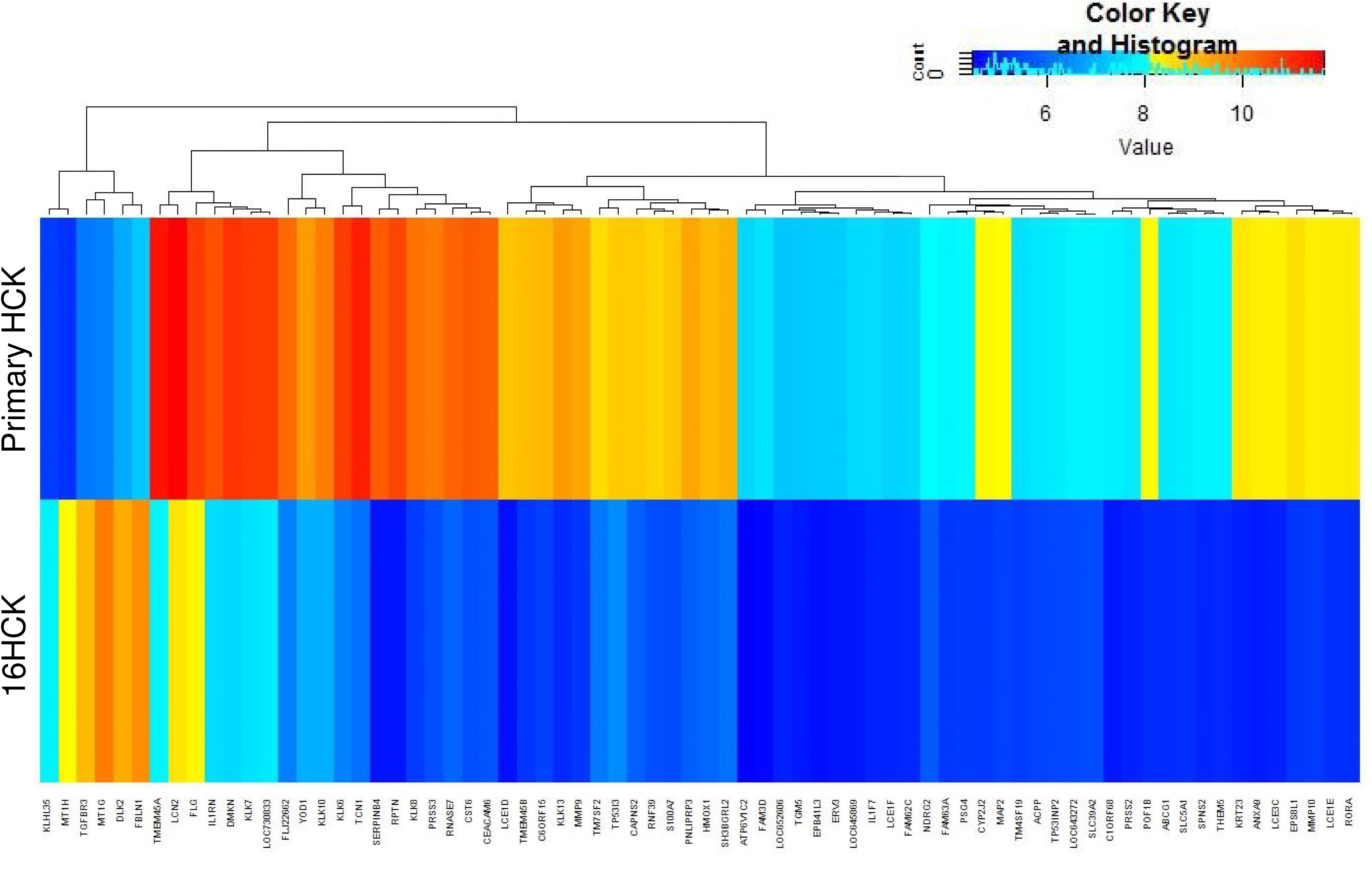
Heat map of significantly modified genes. Raft experiments were set up with three primary HCK and 16HCK cell lines in duplicates, and the tissue were harvested at 10 days of culture for RNA extraction and microarray analysis. The heat map shows genes that were significantly modulated (p < 0.05) at least 5-fold with HPV16 infection.

### Gene ontology analysis

In order to identify biological pathways that are significantly affected with HPV16 infection, we conducted gene ontology (GO) analysis using the online tool GOrilla (29). The GO analysis result was then summarized using REVIGO to combine similar GO terms for simplified visualization (30). 144 GO terms that were significantly represented in upregulated genes were summarized to 82 GO terms (Table S2), and 77 GO terms that were significantly represented in downregulated genes were summarized to 52 GO terms using REVIGO (Table S3).

Amongst the genes that were upregulated with HPV16 infection, many of the represented GO terms are associated with cell cycle progression (cell cycle process, cell division, cell cycle phase transition, regulation of mitotic cell cycle, and cell cycle) and DNA metabolism (DNA metabolic process, DNA repair, cellular response to DNA damage stimulus) as shown in Fig. 2A. This suggests that persistent infection with HPV16 drives cellular proliferation in cervical tissue as expected due to the presence of viral oncogenes E6/E7 and consistent with previous gene expression studies (16, 19, 27, 31). It is known that HPV oncoproteins E6 and E7 inhibit tumor suppressor genes *TP53* and *RB1* respectively, and that this is the main mechanism through which the virus promotes proliferation and tumorigenesis. “Translesion synthesis,” “neuron projection regeneration,” and “retina morphogenesis in camera-type eye” were amongst the upregulated DNA metabolism GO terms which have not yet been reported by previous global gene expression studies of HPV16 infection. Translesion synthesis is a cellular DNA damage tolerance process of recovering from stalled replication forks by allowing DNA replication to bypass certain lesions (32, 33). Eight genes of the translesion synthesis GO category were upregulated in our analysis including *POLE2*, *UBA7*, *MAD2L2*, and *RPA2* which have not yet been reported in previous global gene expression studies of HPV infection. So far, not much is known about the exact role of translesion synthesis in HPV infection. One previous study speculated that HPV oncoprotein E6 may have inhibitory effects on translesion synthesis (34) while another study reported that p80, a cellular cofactor of HPV31 replication, may downregulate translesion synthesis (32). Our study shows for the first time that translesion synthesis is upregulated in a productive raft culture model of HPV16, suggesting that this process may facilitate viral replication and production.

**FIG 2.**
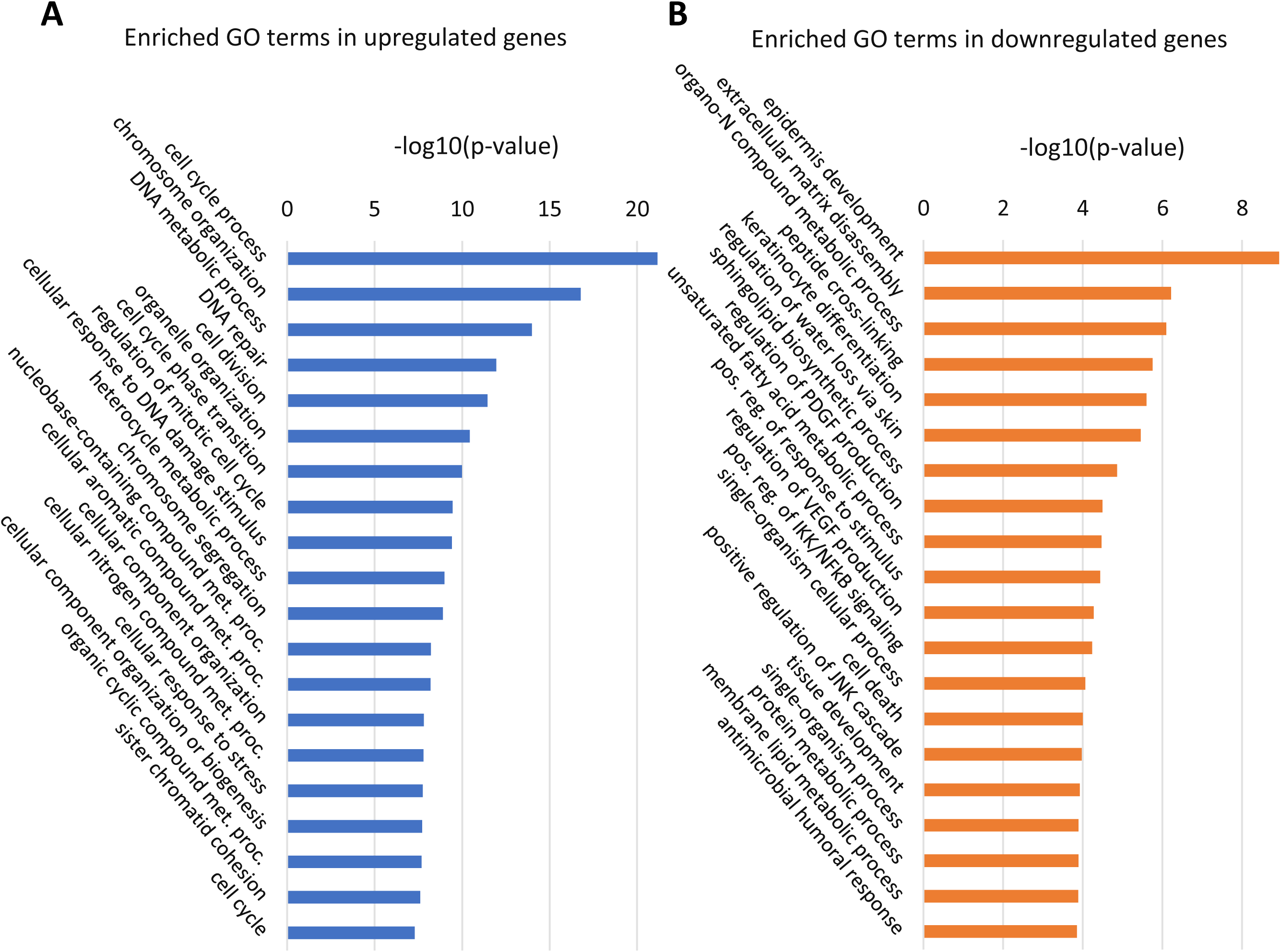
Enriched GO terms with lowest p-values. GOrilla was used to perform initial GO analysis of our microarray data, and REVIGO was used to summarize the enriched GO genesets from the upregulated (A) and downregulated (B) list of genes. The entire list of GO terms can be found in Table S2 and Table S3. Met. proc.: metabolic process.

Within the genes that were downregulated with HPV16 infection, the GO terms that were significantly represented are associated with skin development (epidermis development, keratinocyte differentiation), immune response (positive regulation of IKK/NF-κB signaling, antimicrobial humoral response), and cell death (cell death, positive regulation of JNK cascade) as shown in Fig. 2B. Downregulation of genes associated with skin development have also been observed in previous studies of HPV-positive tumors (13, 17, 35). Similarly, downregulation of genes under GO terms associated with immune response and cell death have been shown in previous studies of HPV-positive tumors and E6 transgenic mice (18, 22, 23, 28, 36, 37). Overall, these results suggest that HPV16 perturbs normal epidermal development while having a hyperproliferative and immunosuppressive effect on cervical tissue. Downregulated GO terms that have not yet been reported in previous global gene profiling studies of HPV infection included “positive regulation of cell migration” and “regulation of platelet-derived growth factor production.” In contrast to our results, previous studies of global gene expression at cancerous stages of HPV infection have shown upregulation of genes involved in cell migration, which is an indicator of invasive, or metastatic cancer (20, 36, 38). Specifically, one study showed that *MMP9*, a well-established metastatic gene, is upregulated in cervical carcinoma cell lines and tissue samples, whereas our microarray analysis shows that this gene is downregulated 17.1-fold (38). Moreover, *LCN2*, which is overexpressed in various cancers and prevents degradation of *MMP9*, was downregulated 9.1-fold in our microarray analysis (39–46). These opposing trends in modulation of cell migration genes highlights the fact that our study investigates early productive stages of HPV infection, whereas the other studies focus on the cancerous stages of infection when viral production is significantly decreased. We speculate that cell migration genes interfere with efficient viral replication and assembly and, therefore, are suppressed during the productive stages of infection, whereas these genes are upregulated to facilitate tumor development and metastasis once the infection enters the cancerous stages.

### Upregulation of cell cycle and DNA metabolism

While GO analysis revealed biological processes that are significantly affected by persistent infection of HPV16, it does not provide information on interactions amongst the genes. Therefore, we further examined the interaction amongst individual proteins based on published data using the online tool STRING (47, 48). STRING protein-protein interaction network analysis allows us to identify signaling pathways, interactions amongst signaling pathways, and central proteins that have maximum interactions with other proteins within a category. A protein association network was created for each of the five aforementioned categories of biological processes that were broadly represented in the GO analysis: cell cycle, DNA metabolism, skin development, immune response, and cell death.

From the upregulated genes, a protein association network was created with genes from 32 GO terms related to cell cycle (Fig. 3A, Table S4). Genes that are not known to be associated with any other gene in this group were excluded from the figure for simplified visualization. The protein association analysis revealed 15 genes that are central to the network as highlighted in red in Fig. 3A. These genes include nucleoporins (*NUP 43, 85, 107*), centromere proteins (*CENP E, F, P*), and kinetochore-associated proteins (*KNTC1, SKA2*). This suggests that HPV infection drives the cell cycle and increases the expression of structural proteins involved in cell cycle and cell division. Amongst these genes, *NUP43*, *NUP85*, *NUP107*, *CENPP*, and *SKA2* have not yet been reported to be associated with HPV infection by previous gene expression studies.

**FIG 3.**
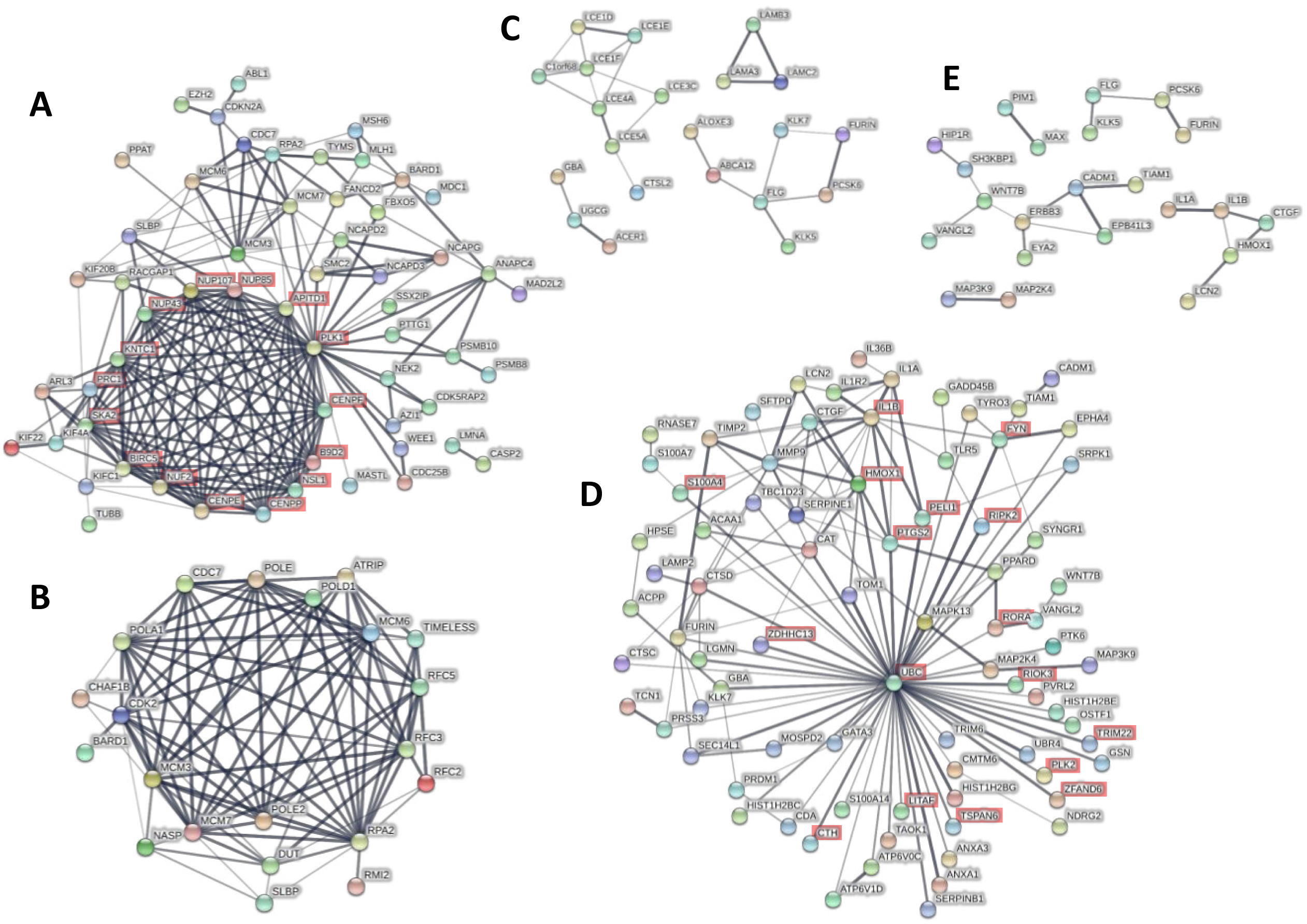
Protein association networks. Online tool STRING was used to create protein functional association networks of upregulated (A-B) and downregulated (C-E) genes with >1.5-fold modulation (p<0.05). (A) Genes from GO categories related to cell cycle regulation were combined to create the network. (B) Genes from GO categories related to DNA metabolism were combined to create the network. (C) Genes from GO categories related to skin development and differentiation were combined to create the network. (D) Genes from GO categories related to inflammation and immune response were combined to create the network. (E) Genes from GO categories related to apoptosis and cell death were combined to create the network. Thicker and darker lines represent greater confidence in protein interaction based on supporting data. Genes that had no connection to any other gene within the network were excluded from the diagram.

Similarly, a protein association network was created with genes from 42 GO terms related to DNA metabolism (Fig. 3B, Table S5). Genes in this network include minichromosome maintenance complex components (*MCM 3, 6, 7*), replication factors (*RFC 2, 3, 5*), and DNA polymerase subunits (*POLA1, POLE, POLE2, POLD1*). Amongst these genes, *POLA1*, *POLE*, and *POLE2* have not yet been reported to be associated with HPV infection by previous gene expression studies. Previous studies have shown that overexpression of MCM genes are correlated to cervical carcinogenesis and specifically, *MCM7* has been shown to interact with HPV18 E6 oncoprotein (49, 50). These results are consistent with upregulation of cell cycle genes since cell division requires DNA replication and proteins associated with DNA metabolism. Additionally, our result suggests that the upregulation of MCM genes is initiated at early stages of HPV infection and sustained throughout carcinogenesis.

### Downregulation of skin development, immune response, and cell death

Within the downregulated gene sets, we combined genes from 7 GO terms in the protein association analysis for skin development, which gave 4 small networks (Fig. 3C, Table S6) including a network of late cornified envelope genes (*LCE 1D, 1E, 1F, 3C, 4A, 5A*) and a network of laminins (*LAMA3*, *LAMB3*, *LAMC2*). All eight genes in the network of LCEs were included in the GO term “epithelial cell differentiation” while the three laminin genes were included in the GO term “epidermis development.” The LCE genes encode stratum corneum proteins of the epidermis, and are all located in the same region of chromosome 1 (1q21.3) suggesting that epigenetic modifications might be suppressing the expression of this region as a whole. Our study shows for the first time the downregulation of a network of LCE genes in HPV infection. The two previous studies that analyzed global gene expression changes in early stage HPV infection may not have observed changes in LCE genes since they used monolayer cultures that do not allow formation of the four layers of differentiating keratinocytes in the epidermis (22, 23). The three laminin genes *LAMA3*, *LAMB3*, and *LAMC2* encode the three subunits that make up laminin 5, which plays an important role in wound healing, keratinocyte adhesion, motility, and proliferation (51, 52).

Genes from 16 GO terms were included in the protein association analysis for inflammation (Fig. 3D, Table S7). Ubiquitin C (*UBC*), which is downregulated 2.48-fold with HPV16 infection, is at the center of this network with the most associations with other genes related to inflammation. *UBC* is one of the two polyubiquitin genes that are involved in various cellular processes including protein degradation, protein trafficking, cell-cycle regulation, DNA repair, and apoptosis (53). Other ubiquitin-related genes that were significantly downregulated in our microarray analysis includes proteins that are involved in ubiquitin conjugation (*UBE2G1*, *UBE2F*, *UBR4*, *UBTD1*), whereas two of the four ubiquitin-related genes that were significantly upregulated are involved in deubiquitination (*USP1*, *USP13*). This suggests that a decrease in ubiquitination may be important in the HPV16 life-cycle, and that the virus is trying to achieve this by decreasing ubiquitination and increasing deubiquitination. In previous studies, UBR4 has been shown to be a cellular target of HPV16 E7 oncoprotein (54, 55), and we have shown that HPV16 upregulates deubiquitinase UCHL1 in order to escape host immunity (56). Since ubiquitins are involved in many cellular processes, it is hard to identify which specific pathway is being affected by the decrease in ubiquitination. It is possible that the virus is preventing degradation of proteins that are targeted by ubiquitin. Also, since our results suggest that HPV infection drives cell cycle and downregulates cell death, it is possible that the downregulation of ubiquitination is involved in these processes. The network also included cytokines (*IL1A*, *IL1B*, *IL36B*), MAP kinases (*MAPK13*, *MAP2K4*, *MAP3K9*), proteases (*CTSD*, *CTSC*, *KLK7*, *FURIN*), serine protease inhibitors (*SERPINB1*, *SERPINE1*), and antimicrobial genes (*S100A7*, *RNASE7*, *PRSS3*, *HIST1H2BC*, *HIST1H2BE*, *HIST1H2BG*, *LCN2*). Amongst these genes, *MAP3K9*, *CTSD*, *CTSC*, *SERPINE1*, *HIST1H2BC*, *HIST1H2BE*, and *HIST1H2BG* have not yet been reported to be associated with HPV infection by previous gene expression studies. Of note, RNASE7 is a broad-spectrum antimicrobial protein that we have previously shown to be downregulated by HPV infection (57). In terms of specific signaling pathways, regulation of I-kappaB kinase (IKK) and NF-κB signaling was significantly represented in the network (Fig. 3D, highlighted in red), which is consistent with our previous study that showed suppression of NF-κB activation by HPV16 (58). Most of these genes were shown to interact with UBC (Fig. 3D), and it is known that ubiquitination and proteolytic degradation of NF-κB inhibitor IκB can lead to NF-κB activation (59). This suggests that HPV16 is evading the immune system by suppressing the NF-κB pathway, and that this suppression may be mediated by downregulation of UBC.

Lastly, genes from 4 GO terms were included in the protein association analysis for cell death and gave 5 small networks (Fig. 3E, Table S8). The networks include genes from the JNK signaling pathway (*TIAM1*, *IL1B*, *VANGL2*, *CTGF*, *WNT7B*) suggesting that HPV16 may be suppressing cell death, and promoting transformation and tumorigenesis through this pathway. This is consistent with our previous study that showed downregulation of cell death with HPV infection (60). Both NF-κB and JNK pathways are downstream of TNF signaling, and TNF ligands and receptors (*TNFRSF19*, *LITAF*, *TNFSF9*) were downregulated in the microarray. This suggests that HPV16 is downregulating NF-κB and JNK pathways via TNF signaling downregulation. Amongst these genes, *TIAM1*, *VANGL2*, *WNT7B*, *TNFRSF19*, and *LITAF* have not yet been reported to be associated with HPV infection by previous gene expression studies.

### Gene transcription changes correlate with changes in protein expression

Of the numerous biological processes that were identified with GO analysis, we wanted to focus on processes and pathways that we felt were unique and most relevant to HPV infection and life-cycle. Therefore, four genes that were modulated at least 10-fold were selected for validation from processes involving epidermal development and differentiation (*KLK8*, *RPTN*, *KRT23*), and immune response (*SERPINB4*). Additionally, *UBC* was selected for validation since it was shown to be central to the protein association network of inflammation (Fig. 3D), and cyclin-dependent kinase 2 (*CDK2*) was included for analysis as a marker of proliferation. In our microarray analysis *KLK8*, *RPTN*, *KRT23*, *SERPINB4*, and *UBC* were downregulated 25.8, 57.5, 11.1, 48, 2.48-fold, respectively, and *CDK2* was upregulated 1.74-fold with HPV16 infection. KRT23 is a structural protein in epithelial cells, whereas KLK8 and RPTN are involved in skin barrier proteolytic cascade and cornified cell envelope formation, respectively (61, 62). Recently, studies have reported that KRT23 may be involved in other cellular processes including cell cycle regulation and apoptosis (63), which are key processes that are modulated by HPV infection in our study. KRT23 has not yet been reported by any other gene expression studies of HPV infection and, therefore, could be developed into a novel biomarker of productive HPV infection. In a recent gene expression profiling study, SERPINB4 was shown to be downregulated in early stage HPV infection consistent with our analysis (23). SERPINB4 is a serine protease inhibitor that is overexpressed in inflammatory skin diseases and various cancers including squamous cell carcinomas, and may play a critical role in the immune response against HPV replication and virion production as it has been shown that increased SERPINB4 expression can activate NF-κB (64–70). Similarly, KLK8 has been shown to be overexpressed in cervical cancer, ovarian cancer, and oral squamous cell carcinoma (71–74). So far, not much is known about these proteins in the context of pre-neoplastic HPV infections, and therefore, they could potentially become biomarkers or therapeutic targets of HPV infection at its early stages. In particular, overexpression of SERPINB4 and KLK8 in cancers prominently contrasts our microarray data that includes the two genes amongst the top downregulated genes. This contrast highlights the different microenvironments of the precancerous and cancerous states of HPV infection, and understanding the role of SERPINB4 and KLK8 may provide critical insight into the mechanism of HPV-induced carcinogenesis. For the validation experiments, six new raft cultures were put up with uninfected keratinocytes isolated from different cervical biopsies and six 16HCK raft cultures were put up from three 16HCK cell lines in duplicates. In order to validate the microarray results in new tissue samples, all of the raft cultures were set up with new cell lines except for one set of 16HCK rafts.

Downregulation of *KLK8*, *RPTN*, *SERPINB4*, and upregulation of *CDK2* at the transcriptional level was observed with RT-qPCR consistent with the microarray analysis (Fig. 4). However, transcription of *UBC* didn’t show statistically significant modulation with HPV16 infection. We then further validated downregulation of KLK8, RPTN, KRT23, and SERPINB4 at the translational level with western blot (Fig. 5A). Although KLK8 expression was visibly downregulated with HPV16 infection in western blot, statistical significance was not reached. To measure UBC protein expression, we used an anti-ubiquitin antibody as a proxy for measuring UBC translation since UBC is simply a polyubiquitin protein that accounts for the majority of basal level ubiquitin in cells (53, 75). Western blot against ubiquitin showed differential ubiquitination of various proteins with HPV16 infection and downregulation of monoubiquitin (Fig. 5B). Lastly, downregulation of RPTN, KRT23, SERPINB4, and Ubiquitin were validated with immunofluorescence staining (Fig. 6). In particular, RPTN, KRT23, and SERPINB4 are minimally expressed in the basal layers of both infected and uninfected controls with no significant difference in expression between the two groups. In contrast, the three proteins are strongly expressed in the upper layers of uninfected tissues and this expression is significantly decreased with HPV16 infection. The downregulation of the proteins may be attributed to the loss of the cornified layer in infected raft tissues.

**FIG 4.**
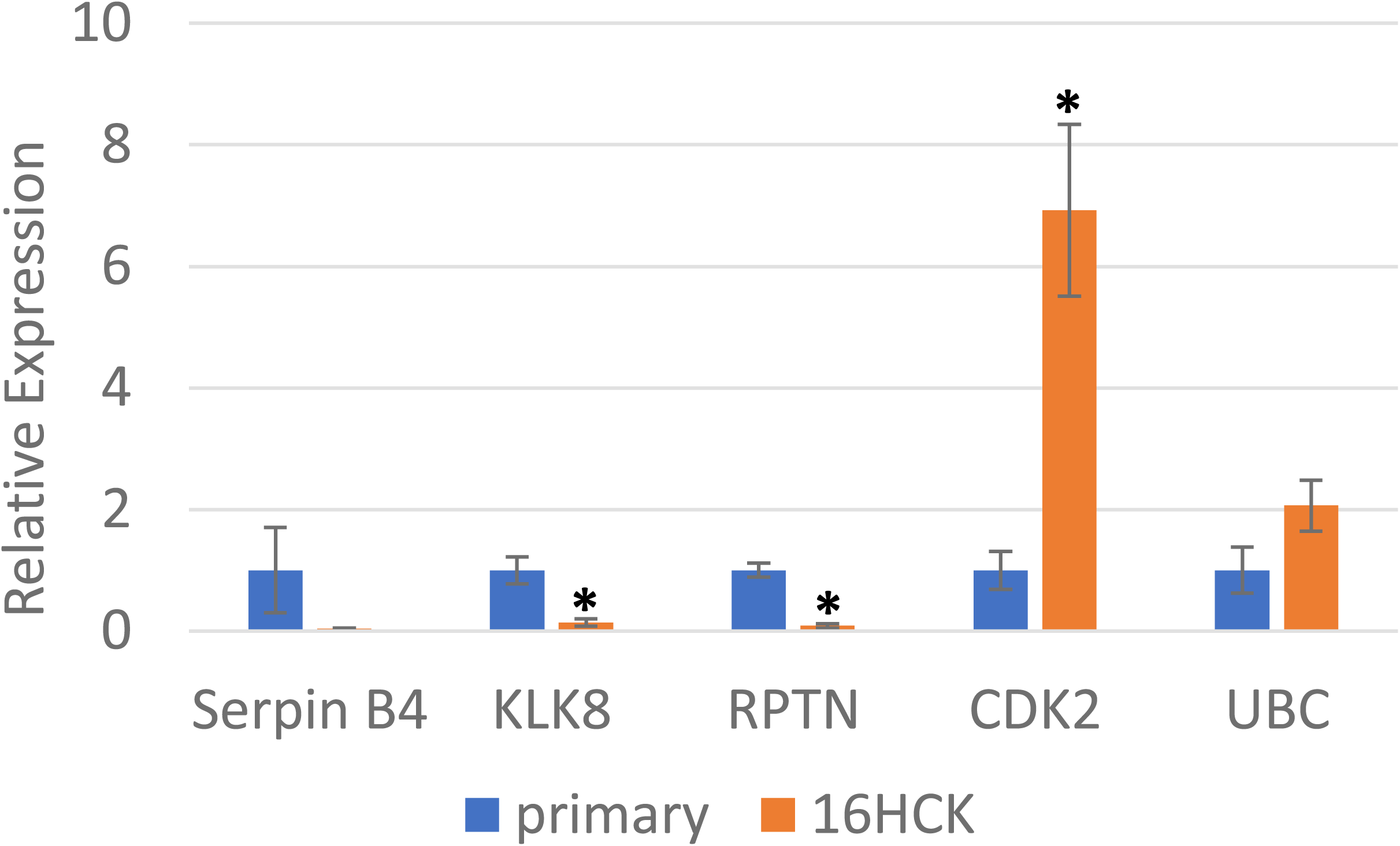
RT-PCR of genes of interest. Raft experiments were set up with three 16HCK cell lines in duplicates and six primary HCK cell lines. The raft tissues were harvested at 20 days of culture. RNA was extracted from the tissues and RT-PCR was performed to measure transcription levels of SERPINB4, KLK8, RPTN, CDK2, and UBC. Transcription levels of TATA-binding protein (TBP) was used as control to normalize each measurement (statistical significance p<0.05).

**FIG 5.**
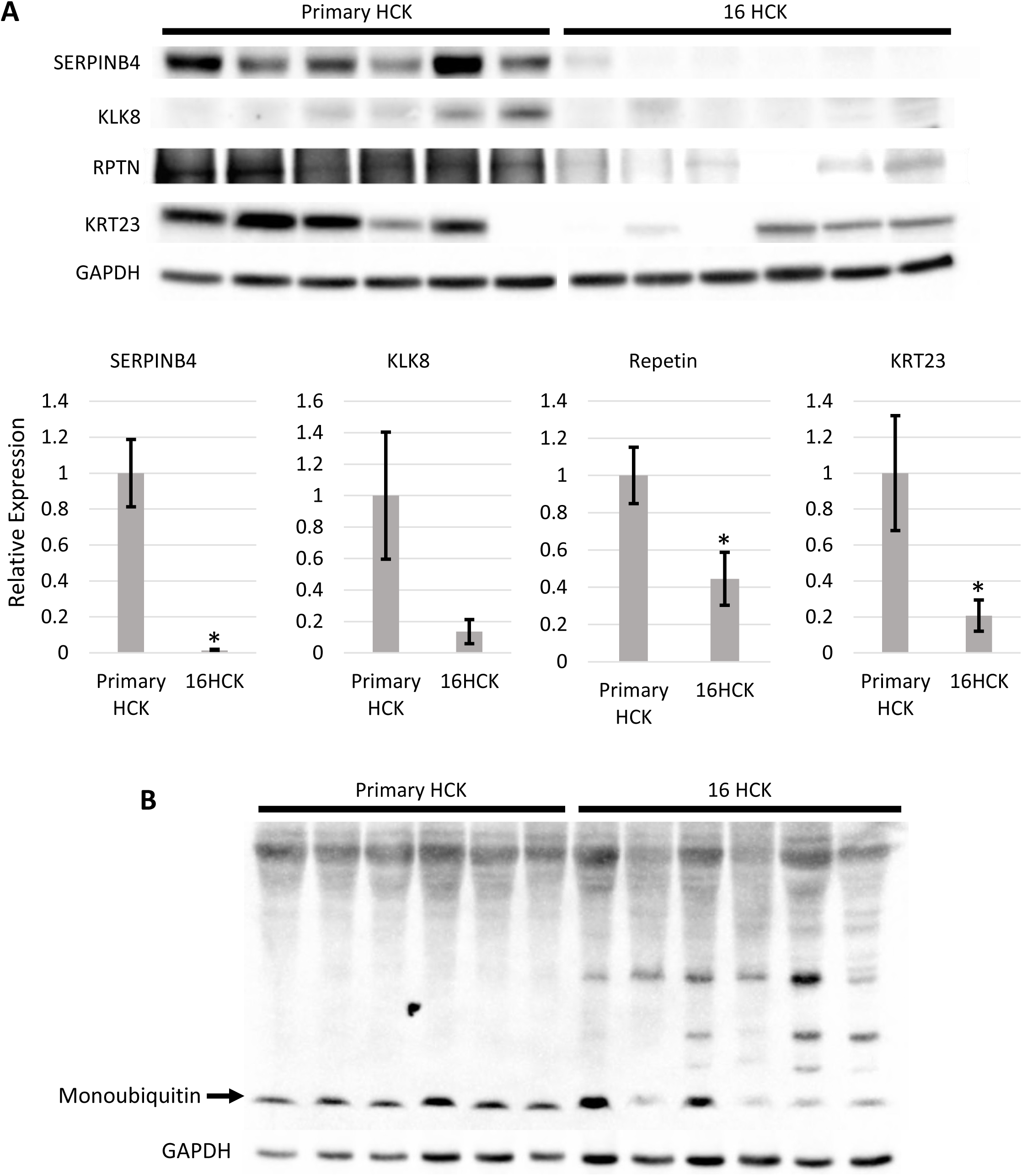
Western blot analysis of raft tissue. (A) Protein expression of SERPINB4, KLK8, RPTN, and KRT23 were tested with Western blots in primary HCK and 16HCK raft tissue harvested at 20 days of culture. Densitometry analysis was conducted on the Western blots (statistical significance p<0.05). (B) Protein expression of Ubiquitin was tested with Western blot in primary HCK and 16HCK raft tissue harvested at 20 days of culture. Experiments were conducted in three 16HCK cell lines in duplicates and six primary HCK cell lines. GAPDH was used as control. Images were acquired with Bio-Rad ChemiDoc MP Imaging System and Image Lab Version 6.0.0 software.

**FIG 6.**
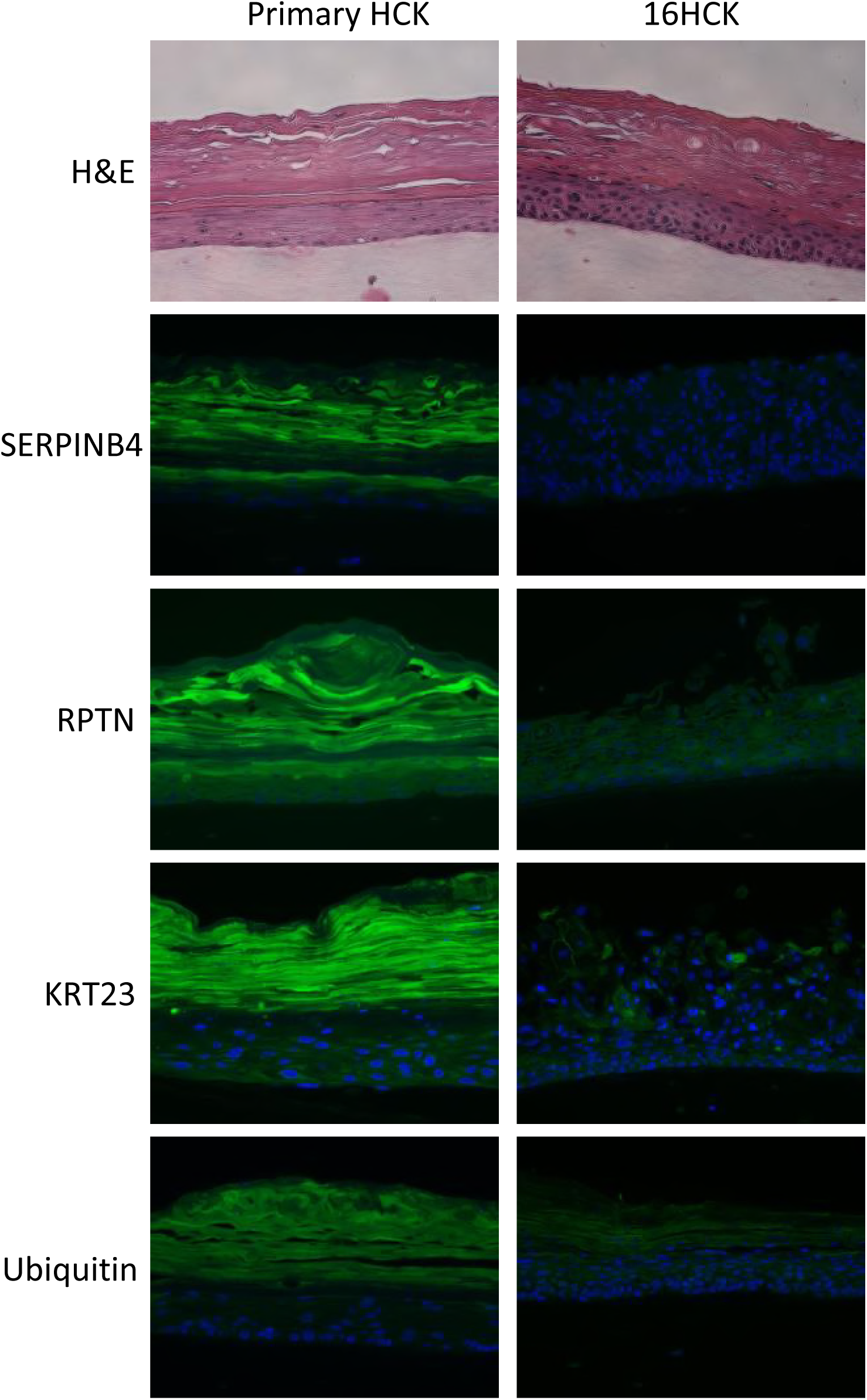
Immunofluorescence staining of raft tissue. The spatial protein expression of SERPINB4, RPTN, KRT23, and Ubiquitin was observed by performing immunofluorescence staining of primary HCK and 16HCK raft tissue fixed at 20 days of culture (green: target protein, blue: nuclear staining). Experiments were conducted in three 16HCK cell lines in duplicates and six primary HCK cell lines. Magnification 200×. H&E: hematoxylin and eosin staining. Images were acquired with Nikon Eclipse 80i microscope and NIS Elements version 4.4 software.

## Discussion

Despite widespread screening for cervical cancer and development of HPV vaccines in recent decades, the burden of cervical cancer remains to be one of the highest amongst female cancers worldwide. In an attempt to understand HPV infection and its progression to cancer at a holistic level, many studies have investigated global gene expression changes that occur with HPV infection. Most of these studies focus on neoplastic lesions and cancerous lesions (13–21). This is because early stages of HPV infection, which typically lasts years, is often cytologically and clinically asymptomatic until it reaches neoplastic stages of infection and, therefore, the majority of clinical samples are limited to these late cancerous stages of infection. As a result, there is a gap in knowledge in global gene expression at pre-neoplastic stages of HPV infection. In two previous studies, early passage HPV16-immortalized human keratinocytes (22) and spontaneously immortalized human keratinocytes transfected with the HPV16 genome (23, 76) were used to measure global gene expression changes at precancerous stages of HPV infection. However, these two studies used keratinocytes derived from foreskin, which is not ideal for modeling HPV infection in cervical tissue since the virus may have tissue-specific effects (24). Moreover, these studies used monolayer cell cultures that do not allow the virus to progress through its entire life-cycle. Amongst the genes that were reported to be modulated in the two studies, only 28.1% and 1.6% of them matched our results, and moreover, some of the genes were modulated in the opposite direction (22, 23). The stark discrepancies between our study and the two previous studies can be attributed to our use of organotypic raft culture system instead of monolayer cell cultures. The HPV life-cycle is dependent on the various stages of keratinocyte differentiation that occur in the epidermis of the skin. Since the HPV life-cycle spans all layers of the epidermis (basale, spinosum, granulosum, corneum), monolayer cell cultures cannot produce progeny virus particles and, therefore, are limited models of HPV infection. In our study, we overcome these limitations by using human cervical keratinocytes and by creating organotypic raft cultures that allow the full progression of the HPV life cycle. Raft cultures were also made from uninfected primary HCKs to serve as control. Viral titers were measured on all 16HCK raft tissues to check for high levels of viral particle production (Table 1), which confirms that the viral genome is maintained episomally allowing productive infection. Integration of viral genome into the host genome, and subsequent reduction in viral particle production is a hallmark event in the progression of precancerous to cancerous lesions and thus, high levels of viral particle production indicates that the infection is in its earlier precancerous stages. Our study presents for the first time global gene expression changes in cervical tissue with productive HPV infection.

Our microarray data showed that the majority of the modulated genes are downregulated (Fig. 1). Gene ontology analysis of the microarray data identified gene categories that were significantly represented including cell cycle progression and DNA metabolism in the upregulated genes, and skin development, immune response, cell death in the downregulated genes (Fig. 2). The upregulation of cell cycle and DNA metabolism genes, and downregulation of cell death genes reflect the proliferative nature of persistent HPV16 infection and is likely the result of viral oncogenes E6 and E7 inhibiting the cell cycle regulatory genes *TP53* and *RB1*, respectively. Downregulation of immune response and skin development genes can be understood in the context of the virus modulating the host environment to achieve efficient replication and virion production. The trends of modulation in these five gene categories are consistent with previously reported studies of global gene expression (13, 16–19, 22, 23, 27, 28, 31, 35–37).

Several genes were selected for validation at the transcription and translational levels based on the degree of fold-change, relevance to the HPV life-cycle, and protein association network analysis. *KLK8* and *RPTN* were selected for validation as they were amongst the top downregulated genes and are both involved in epithelium development. It is not surprising to see many genes involved in epithelium development to be affected by HPV infection since the virus infects, replicates, and assembles in the epithelium. In particular, HPV is not a lytic virus and is released from the epidermis via desquamation. Repetin was downregulated 57.5-fold with HPV16 infection in our microarray analysis and this downregulation was validated with qPCR, western blot, and IF staining. Repetin is a component of the epidermal differentiation complex, and is involved in the formation of the cornified cell envelope (CE) (62). The CE is an insoluble matrix of covalently linked proteins formed beneath the plasma membrane of differentiating keratinocytes and plays an important role in the skin’s function as a protective physical barrier against the external environment (77). In the context of HPV infection, the CE may hinder virion release because of its function as a physical barrier. Additionally, a previous study has shown that CEs of epithelial tissue infected with HPV11 are thinner and more fragile compared to those of healthy tissue (78). Therefore, it can be speculated that downregulation of Repetin by HPV may be a strategy to weaken the CE and increase the efficiency of virion release.

*KLK8* was downregulated 25.8-fold in our microarray analysis, which was validated with qPCR and western blot. *KLK8* was the most significantly downregulated gene of the seven KLK genes that were downregulated with HPV16 infection: *KLK3*, *KLK5*, *KLK6*, *KLK7*, *KLK10*, *KLK13* were downregulated 3.6, 4, 22.1, 8.6, 7.5, 21.5-fold, respectively. A previous gene expression study has also identified a cluster of KLK genes (*KLKs 5, 6, 7, 10, 11*) that are downregulated at early stages of HPV16 infection (22). KLKs are a family of 15 serine proteases that are clustered on chromosome 19q13.4 and one of their main functions is cleaving corneodesmosomal adhesion molecules in the cornified layer of the epidermis, which allows regulated desquamation of keratinocytes (61, 79–81). It is counterintuitive that HPV16 infection downregulates KLKs since the virus is released via desquamation. We speculate that the rate at which normal epithelium desquamates is faster than the rate at which HPV virions mature in the cornified layer, and therefore, the virus may be downregulating KLKs in order to impede desquamation and allow virions to adequately mature before being released to the surrounding environment. *KLK8* also plays a role in activation of the antimicrobial peptide LL-37 and thus, HPV16 may be downregulating the protein in order to prevent antimicrobial reaction (61, 82).

SERPINB4, also known as squamous cell carcinoma antigen 2 (SCCA2), is a member of the serpin family of serine protease inhibitors and was downregulated 48-fold in our microarray analysis. The downregulation in microarray analysis was validated with qPCR, western blot, and IF staining. SERPINB4 along with SERPINB3 (squamous cell carcinoma antigen 2; SCCA1) have been shown to be overexpressed in various types of cancers including cervical, esophageal, lung, breast, and liver cancers (65, 67–69, 83–85). One of the mechanisms through which SERPINB4 contributes to tumor maintenance is the inhibition of granzyme M-induced cell death(86). Additionally, SERPINB4 overexpression is associated with inflammatory diseases including psoriasis and atopic dermatitis (64, 66, 87–89). Remarkably, *KLK8* shares the same expression pattern in these diseases: while our microarray analysis shows significant downregulation during productive HPV16 infection, the overexpression of *KLK8* has also been associated with both squamous cell carcinomas and inflammatory skin diseases, such as psoriasis and atopic dermatitis (71–74, 90). We speculate that the virus downregulates *SERPINB4* and *KLK8* during productive infection as part of a broad effort to dampen the inflammatory response. Of particular note is that the two genes are overexpressed in various cancers while in our study they are amongst the top downregulated genes during productive HPV infection. Additionally, we have identified nine other genes that are significantly downregulated in our microarray analysis, but have shown to be overexpressed or contribute to disease progression in various types of cancers (Table 4). This highlights the fact that early productive stages of HPV infection present a vastly different microenvironment and disease state from the cancerous stages of infection when virion production is significantly decreased. This suggests that KLK8 and SERPINB4 may interfere with the HPV life-cycle or contribute to the immune surveillance against the virus, and therefore, are downregulated during productive infection. In contrast, the two proteins may be necessary for tumor maintenance, and therefore, overexpressed during the cancerous stages of infection. However, since we did not measure levels of expression of these proteins at cancerous stages we cannot definitively compare the protein levels between precancerous and cancerous stages. A direct comparison would require creating organotypic raft cultures with cervical cancer cell lines and measuring SERPINB4 and KLK8 protein levels. In future studies, we aim to investigate the mechanism of *KLK8* and *SERPINB4* downregulation, and how the two genes affect HPV entry, intracellular trafficking, replication, and assembly.

**Table 4.** Differential gene expression between productive HPV16 infection and cancers.

| Gene | Expression fold- | Cancers in which gene expression |
| --- | --- | --- |

|  | **change in**<br>**productive HPV**<br>**infection** | **increases and/or promotes disease**<br>**progression** |
| --- | --- | --- |
| *SERPINB4* | -48.0 | HNSCC (65), cervical (67, 69), esophageal (68), lung (100), liver (85), breast (84) |
| *KLK5* | -3.2 | Bladder (101), ovarian (102), lung (103), breast (104), oral squamous cell carcinoma (OSCC) (73) |
| *KLK6* | -22.1 | Bladder (101), ovarian (102), colorectal (105), head and neck squamous cell carcinoma (HNSCC) (106), pancreatic (107) |
| *KLK8* | -25.8 | Bladder (101), ovarian (102), salivary gland (108), cervical (74), OSCC (73), colorectal (107) |
| *KLK10* | -7.6 | Ovarian (102), OSCC (73), pancreatic (107), colorectal (107) |
| *KLK13* | -21.5 | Lung (109), ovarian (110) |
| *MMP9* | -17.1 | Cervical (38), breast (111), HNSCC (112), prostate (113) |
| *MMP10* | -8.0 | OSCC (114), HNSCC (115), esophageal |
|  |  | (116) |
| *LCN2* | -9.1 | Breast (40, 41), esophageal (42), pancreatic (43), ovarian (44), colorectal (45), thyroid (46) |
| *CTSD* | -1.5 | Breast (117), ovarian (118) |
| *CTSV* | -2.8 | Breast (119), colorectal (119) |

UBC was the central protein in the network of inflammatory genes that were significantly downregulated with persistent HPV16 infection (Fig. 3D). Although qPCR did not show a statistically significant modulation of *UBC* expression, western blot of ubiquitin showed downregulation of monoubiquitin and differential pattern of protein ubiquitination while IF staining showed downregulation of the protein with persistent infection (Fig. 5, Fig. 6). Since ubiquitin is involved in numerous biological processes it is hard to conclude which specific pathways are affected with HPV16 infection. However, many proteins that are associated with UBC in the protein association network (Fig. 3D) are a part of the NFKB pathway, and we speculate that UBC plays a central role in downregulating this pathway especially since it has been shown that ubiquitin-proteasome pathway plays a role in NF-κB activation. In future studies, we would like to investigate the role of *UBC* in relation to top downregulated genes associated with inflammation, such as *SERPINB4* and *KLK8*.

In conclusion, our study shows for the first time global gene expression changes with productive HPV16 infection in an organotypic raft culture model. With gene ontology analysis, broad gene categories were identified that were significantly modulated with persistent HPV16 infection, and these results were largely consistent with what previous studies have reported. In particular, we identified top downregulated genes that have not yet been extensively studied in the context of HPV infection, and that have potential to be developed as therapeutic targets or biomarkers. Moreover, expression patterns of *SERPINB4* and *KLK8* highlighted that precancerous and cancerous stages of HPV infection are two distinct disease states. Although some of the observed gene modulations were consistent with what was expected from results of previous studies, we also observed novel changes that have not been reported before. We attribute these new findings to our unique model of organotypic raft cultures at early stage HPV16 infection, which allows the virus to go through its complete life-cycle. Future studies investigating how the regulation of *SERPINB4* and *KLK8* changes throughout the different stages of infection may shed light on unidentified mechanisms of HPV persistence and tumorigenesis.

## Materials and Methods

### Creating cervical cell lines and organotypic raft cultures

Primary human cervical keratinocytes (HCK) were isolated from cervical biopsies as previously described (91). The Human Subjects Protection Office of the Institutional Review Board at Penn State University College of Medicine screened our study design for exempt status according to the policies of this institution and the provisions of applicable federal regulations and, as submitted, found not to require formal IRB review because no human participants are involved as defined by the federal regulations.

HCK cell lines persistently infected with HPV16 (16HCK) were produced by electroporating primary HCK with HPV16 plasmid DNA as previously described (92, 93). The electroporated cells were cultured with mitomycin-C-treated (Enzo Life Sciences) J2 3T3 feeder cells as previously described (91).

Organotypic raft cultures were grown as previously described (93, 94) at first or second passage for primary HCK and sixth to ninth passage for 16HCK. The raft tissues were harvested after 10 days for microarray analysis and 20 days for qPCR, Western blot, and immunofluorescence staining. Viral gene expression has been shown to peak between 10 and 12 days (95) while virion maturity reaches maximum stability around 20 days (96).

### Microarray analysis

Raft tissue from primary HCK and 16HCK were harvested at 10 days and RNA was extracted using the RNeasy Fibrous Tissue Midi Kit (Qiagen). The experiment was conducted with three primary HCK and three 16HCK samples in duplicates. Microarray analysis was performed using the Illumina HumanHT-12 v4 Expression Beadchip (Illumina, San Diego, CA), which targets over 31,000 annotated genes with more than 47,000 probes derived from the National Center for Biotechnology Information (NCBI) RefSeq Release 38 (November 7, 2009) and other sources. RNA quality and concentration were assessed using an Agilent 2100 Bioanalyzer with RNA Nano LabChip (Agilent, Santa Clara, CA). cRNA was synthesized by TotalPrep Amplification (Ambion, Austin, TX) from 500 ng of RNA according to manufacturer’s instructions. T7 oligo (dT) primed reverse transcription was used to produce first strand cDNA. cDNA then underwent second strand synthesis and RNA degradation by DNA Polymerase and RNase H, followed by filtration clean up. In vitro transcription (IVT) was employed to generate multiple copies of biotinylated cRNA. The labeled cRNA was purified using filtration, quantified by NanoDrop, and volume-adjusted for a total of 750 ng/sample. Samples were fragmented, and denatured before hybridization for 18 hours at 58°C. Following hybridization, the beadchips were washed and fluorescently labeled. Beadchips were scanned with a BeadArray Reader (Illumina, San Diego, CA).

The CLC Genomics Workbench 4.8 package (https://www.qiagenbioinformatics.com/) was used to determine the significantly differentially expressed genes of the HPV16+ versus primary tissue. For each comparison, quantile normalization was performed followed by pairwise homogeneous t-test resulting in normalized fold changes and p-values. Significantly differentially expressed genes were considered to be those with p < 0.05 and absolute fold change >= 1.5.

### Gene ontology analysis and protein association network

In order to categorize the significant gene expression changes into gene ontology (GO) groups the GOrilla package was used (http://cbl-gorilla.cs.technion.ac.il/) (29). Two unranked lists of genes, target (significantly modulated genes) and background (all genes in the microarray), were used to identify significantly enriched GO terms. We focused on the sub-ontology Biological Processes for our analysis. REVIGO was used to further summarize the redundancy in the GO analysis (http://revigo.irb.hr/) (30). In our analysis, we used the similarity coefficient of 0.7 (medium size list) to summarize the GO list.

In order to identify protein-protein associations amongst the upregulated and downregulated genes, the online tool STRING (https://string-db.org/) was used (48). Genes from similar GO categories were pooled together to form protein association networks of “cell cycle” and “DNA metabolism” for the upregulated genes, and “skin development,” “immune response,” and “cell death” for the downregulated genes.

### Viral Titers (DNA encapsidation assay)

Viral titers of each raft experiment were measured with the qPCR-based DNA encapsidation assay as previously described (96, 97).

### RT-qPCR

RT-qPCR was used in order to measure levels of transcription of SERPINB4, KLK8, RPTN, CDK2, and UBC. The experiment was conducted with three primary HCK and three 16HCK samples in duplicates. For*SERPINB4*, forward primer 5’-ATTTCCTGATGGGACTATTGGCAATG-3’, reverse primer 5’-CAGCAGCACAATCATGCTTAGA-3’, and probe 5’-/56-FAM/ACGACACTG/ZEN/GTTCTTGTGAACGCA/3IABkFQ/-3’ was used. For*KLK8*, forward primer 5’-TGGGTCCGAATCAGTAGGT-3’, reverse primer 5’-GCAGGAACATCCACGTCTT-3’, and probe 5’-/56-FAM/CCCTGGATT/ZEN/CTGGAAGACCTCACC/3IABkFQ/-3’ was used. For*RPTN*, forward primer 5’-CCACAAATATGCCAAAGGGAATG-3’, reverse primer 5’-GTCATTTGGTCTCTGGAGGATG-3’, and probe 5’-/56-FAM/ACTGCTCTT/ZEN/GGCTGAGTTTGGAGA/3IABkFQ/-3’ was used. For*CDK2*, forward primer 5’-GCCTGATTACAAGCCAAGTTTC-3’, reverse primer 5’-CGCTTGTTAGGGTCGTAGTG-3’, and probe 5’-/56-FAM/AGATGGACG/ZEN/GAGCTTGTTATCGCA/3IABkFQ/-3’ was used. For*UBC*, forward primer 5’-GGATTTGGGTCGCAGTTCTT-3’, reverse primer 5’-TGGATCTTTGCCTTGACATTCT-3’, and probe 5’-/56-FAM/AGGTTGAGC/ZEN/CCAGTGACACCATC/3IABkFQ/-3’ was used. TATA-binding protein (TBP) was used as control for which forward primer 5’-CACGGCACTGATTTTCAGTTCT-3’, reverse primer 5’-TTCTTGC TGCCAGTCTGGACT-3’, and probe 5’-HEX-TGTGCACAGGAGCCAAGAGTGAAGA-BHQ-1-3’ was used. All primers and probes were synthesized by Integrated DNA Technologies, and QuantiTect Probe RT-PCR Kit (Qiagen) was used for the PCR reactions. All RT-PCR reactions were performed using the C1000 Thermal Cycler (Bio-Rad). The thermal cycler was programmed for 30 minutes at 50 °C, then 15 minutes at 95 °C, then 42 cycles of 15 seconds at 94 °C and 1 minute at 54.5 °C.

### Western blot

Raft tissue were harvested at 20 days and used to prepare total protein extracts as previously described (98). The experiment was conducted with three primary HCK and three 16HCK samples in duplicates. Total protein concentrations were measured using the Peterson protein assay as previously described (99). The total protein extracts were applied to sodium dodecyl sulfate polyacrylamide gel (8-10%) and transferred to nitrocellulose membrane, then incubated overnight at 4 °C with antibodies against SERPINB4 (Lifespan Biosciences, LS-C172681, 1:2000 dilution), KLK8 (Abnova, H00011202-M01, 1:2000 dilution), RPTN (Lifespan Biosciences, LS-B17, 1:2000 dilution), KRT23 (Abcam, ab117590, 1:2000 dilution), and ubiquitin (Cell Signaling, 3933S, 1:2000 dilution). GAPDH antibody (Santa Cruz, sc-47724, 1:1000 dilution) was used as control. The membranes were then washed and incubated with horseradish peroxidase-linked secondary antibody (GE Healthcare, NA931VS/NA934VS) and developed using Amersham ECL Prime Western Blotting Detection Reagent (GE Healthcare). Densitometry analysis was conducted by normalizing the protein expression levels to GAPDH.

### Immunohistochemistry and Immunofluorescence Staining

Raft cultures were grown for 20 days and fixed in 10% buffered formalin, embedded in paraffin, and 4-μm cross-sections were prepared. A section from each sample was stained with hematoxylin and eosin as previously described (26).

For Immunofluorescence staining, the slides were submerged in xylenes for deparaffinization, and then were rehydrated. Antigen retrieval was achieved by submerging the slides in Tris-EDTA buffer (pH 9) in a 90 °C water bath for 10 minutes. The slides were then rinsed with TBS-Tween and blocked with Background Sniper (Biocare Medical). The slides were then stained with the primary antibody overnight at 4 °C. Each sample was stained with antibodies against SERPINB4 (Lifespan Biosciences, LS-C172681, 1:2000 dilution), RPTN (Lifespan Biosciences, LS-B17, 1:1000 dilution), KRT23 (Abcam, ab117590, 1:500 dilution), and Ub (Cell Signaling, 3933S, 1:750 dilution). The slides were then rinsed with TBS-Tween 3 times and stained with secondary antibody (Life Technologies, Alexa Fluor 488) diluted 1:200 for 1 hour at room temperature. Next, the slides were stained with Hoechst nuclear stain (1:5000 dilution) for 15 minutes and rinsed with TBS-Tween twice. All antibodies were diluted in Da Vinci Green diluent (Biocare Medical). The experiment was conducted with three primary HCK and three 16HCK samples in duplicates. A Nikon Eclipse 80i microscope and NIS Elements version 4.4 software was used to acquire images.

### Statistical analysis

In order to establish statistical significance in qPCR data and Western blot densitometry analysis, t-test was used with a p-value cutoff of p < 0.05.

### Accession number

The GEO accession number for our microarray data is GSE109039.

## Acknowledgments

We would like to thank Lynn Budgeon for doing the raft tissue embedding, slicing of sections, and staining.

## Supplemental Material

**Table S1:** Genes significantly modulated with HPV16 infection in human cervical tissue.

**Table S2:** GO analysis of genes upregulated with HPV16 infection in human cervical epithelium. GO analysis performed with GOrilla and summarized with REVIGO.

**Table S3:** GO analysis of genes downregulated with HPV16 infection in human cervical epithelium. GO analysis performed with GOrilla and summarized with REVIGO.

**Table S4:** Upregulated genes: GO terms related to cell cycle.

**Table S5:** Upregulated genes: GO terms related to DNA metabolism.

**Table S6:** Downregulated genes: GO terms related to skin development.

**Table S7:** Downregulated genes: GO terms related to inflammation.

**Table S8:** Downregulated genes: GO terms related to cell death.

